# Transcriptomic analyses of differentially expressed genes, micro RNAs and long-non-coding RNAs in severe, symptomatic and asymptomatic malaria infection

**DOI:** 10.1101/2022.10.27.514083

**Authors:** Mary A. Oboh, Olanrewaju B. Morenikeji, Olusola Ojurongbe, Bolaji N. Thomas

## Abstract

**Background:** Malaria transmission and endemicity in Africa remains hugely disproportionate compared to the rest of the world. The complex life cycle of *P. falciparum* (*Pf*) between the vertebrate human host and the anopheline vector results in differential expression of genes within and between hosts. An in-depth understanding of *Pf* interaction with various human genes through regulatory elements will pave way for identification of additional tool in the arsenal for malaria control. Therefore, the regulatory elements (REs) involved in the over- or under-expression of various host immune genes hold a key to alternative control measures that can be applied for prompt diagnosis and treatment.

**Methods:** We carried out an RNAseq analysis to identify differentially expressed genes and network analysis of non-coding RNAs and target genes associated with immune response in individuals with different clinical outcomes. Raw RNAseq datasets, retrieved for analyses include individuals with severe (Gambia - 20), symptomatic (Burkina Faso - 15), asymptomatic (Mali - 16) malaria as well as uninfected controls (Tanzania - 20; Mali - 36).

**Results:** Of the total 107 datasets retrieved, we identified 5534 differentially expressed genes (DEGs) among disease and control groups. A peculiar pattern of DEGs was observed, with individuals presenting with severe/symptomatic malaria having the highest and most diverse upregulated genes, while a reverse phenomenon was recorded among the asymptomatic and uninfected individuals. In addition, we identified 141 differentially expressed (DE) miRNA, of which 78 and 63 were upregulated and downregulated respectively. Interactome analysis revealed a moderate interaction between DEGs and miRNAs. Of all identified miRNA, five were unique (hsa-mir-32, hsa-mir-25, hsa-mir-221, hsa-mir-29 and hsa-mir-148) because of their connectivity to several genes, including hsa-mir-221 connected to 16 genes. Six-hundred and eight DE lncRNA were identified, including SLC7A11, LINC01524 among the upregulated ones.

**Conclusion:** Our study provides important insights into host immune genes undergoing differential expression under different malaria conditions. It also identified unique miRNAs and lncRNAs that modify and/or regulate the expression of various immune genes. These regulatory elements, we surmise have the potential to serve a diagnostic purpose in discriminating between individuals with severe/symptomatic malaria and those with asymptomatic infection or uninfected.

## Introduction

Malaria remains a huge cause of morbidity and mortality in many endemic regions burdened with the infection especially sub-Saharan Africa (WHO, 2021). The recent World Health Malaria Report estimated about 241 million malaria cases and 627,000 deaths, and 95% of the cases and deaths are from sub-Saharan Africa (WHO, 2021). As a result of the growing threat of parasite and vector resistance to antimalarial drug and insecticides (Abruquah et al., 2010; Adams et al., 2018; Alam et al., 2011; Awolola et al., 2018; Diallo et al., 2022; Fukuda et al., 2021; Hawkes et al., 2015; Oboh et al., 2018; Patel et al., 2017; Straimer et al., 2022; Tacoli et al., 2016; Traoré et al., 2019; Tumwebaze et al., 2021), attention is now focused, more than ever on other control measures that can be deployed for malaria control. Unfortunately, vaccine development has also not been spared from parasite evasive mechanisms. Antigenic variations of different *Plasmodium falciparum* antigens (Abdolaziz Gharaei, 2014; Naung et al., 2022; Pirahmadi et al., 2018) are huge challenges that impede efficacy of the available malaria vaccine. The only approved malaria vaccine gives ~30% protection against malaria with multiple boosts (Gosling and von Seidlein, 2016; Mvi and Gsk, 2015; The Lancet, 2021), thus requiring deeper understanding of the parasite as well as appropriate host responses.

*P. falciparum* parasite possesses a unique complex life cycle criss-crossing the human host, the invertebrate mosquito vector and the pathogen (Mala et al., 2016; Wiser, 2009). Advancement of this life cycle in the different host involves dynamic expression of genes involved in the parasite and the human immune response (Adukpo et al., 2022; Beeson et al., 2016; Chappell et al., 2020; Chen et al., 2014; Cowman et al., 2012; Crompton et al., 2014; Douglas et al., 2011; Duah et al., 2010; Ouédraogo et al., 2011; Tran et al., 2016). Hence, understanding the functional interaction between humans and the parasite becomes pertinent to clarifying insights into the host-pathogen interactome. The genomic architecture of the human host and parasite provides the unique platform shaping host-pathogen interaction, and this communication is a selective force on the host immune response and parasite evasive tactics (Awasthi and Das, 2013; Basu et al., 2013; Duffy et al., 2018; Oboh et al., 2022). Therefore, understanding this interaction require an in-depth analysis of host-pathogen transcriptomic profile. Many studies have been designed to understand parasite transcriptome/genome (Amambua-Ngwa et al., 2019; Tran et al., 2016), but not much work has been done to decipher the transcriptome of the human host under an antigenic assault. Current findings have shown the role of signaling and pathogen recognition receptors in immune response to disease, however, the role of regulatory elements especially in *Pf* infection is currently unknown. A recent report concluded that epigenetically reprogrammed monocytes potentially drive a regulatory disease IL-10, CD163 and CD206 phenotype (Guha et al., 2021), but the regulatory elements involved in the over- or under-expression of these genes are yet to be fully understood, presenting an opportunity to explore a potential area for initiating prevention/ control of this disease.

At present, there are increasing reports of the involvement of regulatory players in modulating the expression of genes at the transcriptional, posttranscriptional, translational and post-translation levels (Sudmant et al., 2015; Xu et al., 2022; Zhai et al., 2020) in different species and diseases state (Morenikeji et al., 2020; Tucker et al., 2021). Such regulatory players include long non-coding RNAs, micro RNAs and transcription factors (TF). LncRNAs are a class of non-coding transcripts, defined by a threshold of >200 nucleotides and found within or between coding genes (Simantov et al., 2022). They have been implicated in regulating different pathological and biological functions at the transcriptional and epigenetic levels, and as such influence host immune response (Morenikeji et al., 2021). MiRNA on the other hand are shorter than lncNRAs, and attached to the 3’-5’ untranslated region and the coding arm of a messenger RNA to modify gene expression, and ultimately protein products translated from modified genes (Simantov et al., 2022). Early diagnosis of infection and quick intervention are key strategies in malaria control. With an array of miRNA and lncRNA discoveries serving as biomarkers in various disease conditions such as cardiovascular diseases, cancer, diabetes and even infectious pathogens (Morenikeji et al., 2019; Wang et al., 2017; Yua et al., 2015), there is an increased opportunity to add these regulatory elements (REs) to the arsenal for malaria control. Many studies on malaria have attempted to correlate differential expression of these REs in individuals with different clinical manifestations to identify host biomarkers that can serve either diagnostic or therapeutic purpose, but none has been found yet.

With an abundance of binding sites for miRNA and mRNA, lncRNA can act as ceRNA (competing endogenous RNA) and are significant regulatory elements in post-transcriptional gene expression (Carpenter et al., 2014; Wissink et al., 2016), while TFs are proteins capable of altering or activating gene-expression level (Boija et al., 2018; Edginton-White and Bonifer, 2022; Mitsis et al., 2020; Sharov et al., 2022). Multiple miRNA databases such as miRWalk (Sticht et al., 2018), miRNet (Chang et al., 2020), and TargetScan (Agarwal et al., 2015) compute potential miRNA-mRNA interactions, while the role of individual miRNA can be inferred through functional analysis with Gene Ontology (GO) (Carbon et al., 2019). Similarly, lncRNA prediction binding software including NONCODE (Bu et al., 2012), lncRNA2Target (Jiang et al., 2015) and lncTar (Li et al., 2014) have been useful in guiding bench experiments. Current algorithms, relying on base pair complementation, evolutionary conservation, and thermodynamic stability of binding regions, have been shown to be useful in predicting miRNA, lncRNA and TF binding sites on target genes (Carpenter et al., 2014; Popp et al., 2021; Wissink et al., 2016).

Therefore, we set out to carry out transcriptomic profiling to identify differentially expressed genes in individuals with different malaria clinical conditions, identify lncRNAs and miRNA and how these interact with immune genes

## Methodology

### RNAseq data acquisition

A total of 107 raw fastq of human RNAseq datasets were retrieved from the Sequence Read Archive (SRA) database. The RNAseq datasets were obtained from individuals infected with severe malaria (The Gambia; n=20), symptomatic (Burkina Faso; n=15), and asymptomatic disease (Mali; n=16), in addition to uninfected controls (Tanzania; n=20; and Mali; n=36). All libraries were paired-end reads with an average of 103 million reads (Table 1); RNAseq reads ranging from 21 - 103 million reads. Data from studies that were not properly defined in terms of infection status at the time of sample collection or for which status of malaria infection (uninfected, asymptomatic, symptomatic and severe) was unknown were excluded from this analysis.

**Table 1:** Countries, accession numbers and infection status of retrieved RNAseq data

### Gene expression analysis of RNAseq data

Low quality reads and any remaining adapters were trimmed off the RNAseq data, using Trimmomatic version 0.38 (Bolger et al., 2014). Hisat2 version 2.2.1 was used to align quality trimmed reads with the human (USCU hg38) reference genome (Kim et al., 2015). The resulting bam files were used as input for constructing the gene counts using htseq-count version 0.9.1 (Anders et al., 2015) with the unstranded option and the annotated hg38 gene transfer format. Gene expression analysis of generated gene counts mapping to the human genome was performed using DESeq2 (version 1.34.0) (Love et al., 2014). Resulting p-value were adjusted using the Benjamin and Hochberg’s approach for controlling false discovery rate. Genes with an adjusted p-value of <0.05 found by DESeq2 were noted as differentially expressed genes (DEGs) involved in human response to the clinical manifestations of malaria. Furthermore, gene counts were filtered and normalized, and low quality expression genes, defined as genes with < 0.5 count per million across all samples, were removed. Samples were clustered by the status of infection (severe, symptomatic, asymptomatic and uninfected) and DEGs were visualized in a heatmap.

### Functional and pathway analysis of human DEGs in clinical manifestations of malaria

Functional enrichment analysis of the topmost differentially expressed genes to determine the molecular functions, biological processes and cellular components associated with these genes was performed using Database for Annotation, Visualization and Integrated Discovery (DAVID). With a false discovery rate (FDR) of 0.1, and a p-value of ≤ 0.05, molecular functions, biological processes and cellular components were catalogued. Generated pathways were further confirmed using the iDEP (Ge et al., 2018) without altering the parameters. KEGG programming algorithm was used to generate significant pathways with a p-value of ≤ 0.05 and FDR of 0.1.

### Differentially expressed miRNA prediction in different disease manifestations

miRNAs that were differentially expressed with a p-value of <0.05 and log fold change ≥ 2 were taken to be upregulated while those with fold change of ≤ −2 and a p-value of <0.05 were downregulated. Some selected miRNAs were used to probe for genes and long non-coding RNA regions on the human reference genome that bind to the 5’-UTR, CDS and 3’-UTR, using miRnet (Chang et al., 2020), TargetScan (Agarwal et al., 2015) and miRTarBase (Huang et al., 2020). Network of the miRNAs interaction with targeted genes and lncRNAs were constructed separately using the miRnet web-based platform (Chang et al., 2020). Additionally, functional enrichment of the differentially expressed miRNAs was carried out using DAVID to determine the molecular functions, biological processes and cellular components connected with these miRNAs.

### Identification of IncRNAs in disease populations

Putative lncRNAs that might potentially impact the expression of immune genes in individuals leading to varied disease manifestation were sorted among the various differentially expressed products. To further locate the lncRNAs on the genome, the NONCODE database, which is a robust repository for non-coding RNAs was used (Bu et al., 2012). Due to the huge size of non-coding RNAs in most vertebrate genomes, some assumptions were made to filter out any non-coding RNAs that might have even the slightest potential of coding for a particular protein. These assumptions include a). lncRNA with more than two exons were selected based on the presumption that high quality transcripts have high number of exons; b). transcripts length of ≥ 200 bp were selected; c). non-coding potential prediction score using the coding-non-coding index (CNCI) of < 0 was used to discriminate between lncRNAs with the ability to code from those without such ability (Guo et al., 2019; Zhang et al., 2017). All lncRNAs that met the above three criteria were selected and a second validation done using the LGC toolkit found among the suite of LnCBook package (Ma et al., 2019) that characterize and identify features of lncRNA, including transcript length. The LnCbook collates lncRNAs from both pre-existing databases and experimentally verified community-curated transcripts making it a reliable database for determining the coding potential of lncRNAs. LncRNAs that were identified to have coding potential by this toolkit were further dropped from the list.

## Results

### Differentially expressed genes in individuals with various malaria clinical manifestations

Of the 107 total RNAseq datasets retrieved from the SRA database and analyzed, 5534 genes were found to be differentially expressed among those with severe, symptomatic, asymptomatic malaria and uninfected individuals. A unique pattern of expression was observed among the different clinical definitions of disease: individuals with severe and symptomatic malaria from Gambia and Burkina Faso had similar expression pattern, with similar genes observed to be upregulated (e.g. UBB, TENTSC, HSP90AA1, FCGR3B, CYBB, SOD2 etc) and downregulated (DCAF12, IFITM3, R3HDM4, CNN2, CFAP65, ATP283 etc). However, a reverse pattern was observed among individuals with asymptomatic malaria infection or uninfected controls. Of these, 3928 genes were upregulated in individuals with severe and symptomatic malaria, and these genes were found downregulated in those with asymptomatic malaria or uninfected individuals. Consequently, 1604 were observed to be upregulated in the asymptomatic or uninfected individuals while being downregulated in the severe/symptomatic groups (Figure 2). One hundred and thirty genes showed no change in their level of expression (Figure 1). Network interactome of the identified genes with various miRNA targets revealed a moderate form (maximum of 4 genes connecting with a miRNA) of interaction where KMT2B, IDF2, TTYH3, NTRK2 genes were observed to interact with hsa-mir-331, EGFR, MBD6, ZNF385A, TTYH3 genes interacted with hsa-mir-24 (Figure 3).

**Figure 1:**
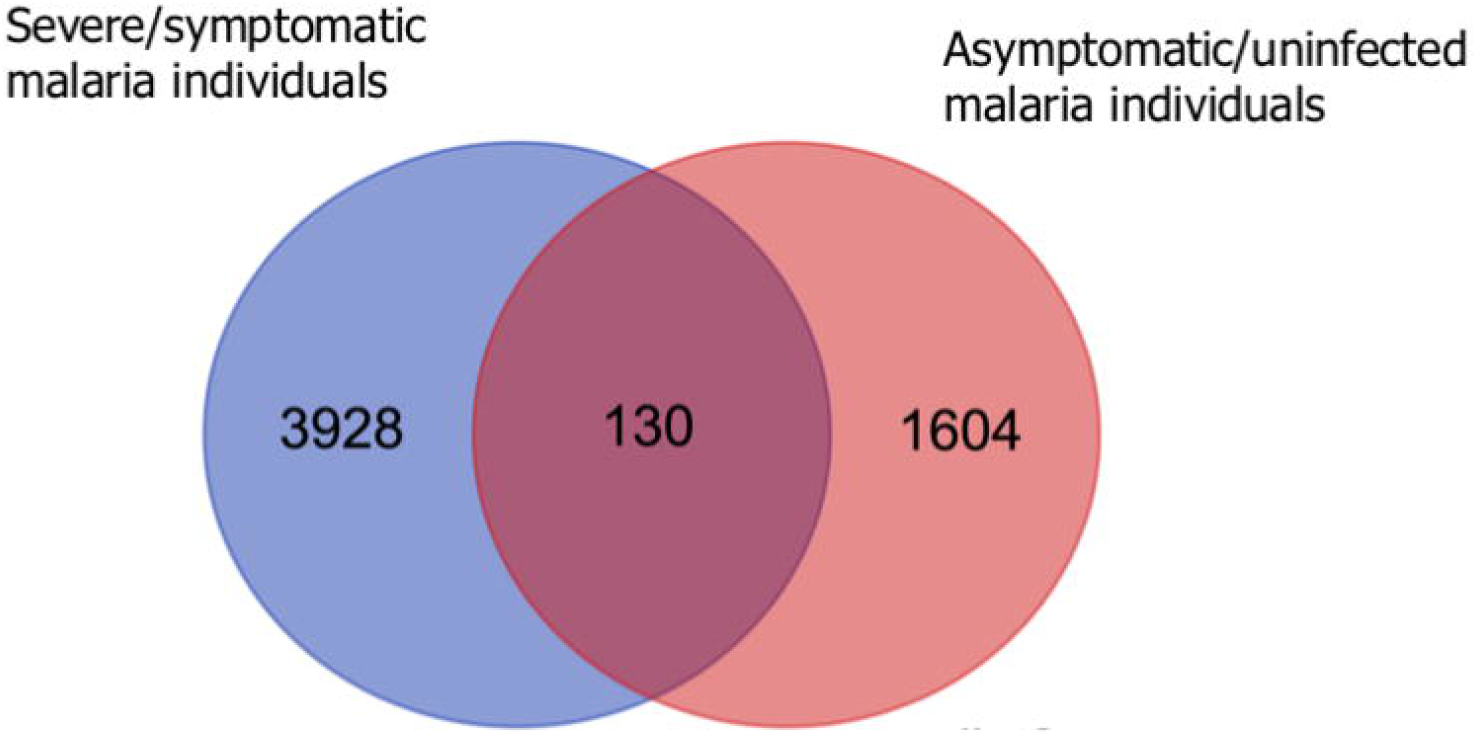
Venn diagram of DEGs in different malaria clinical manifestations. NB: Genes that were upregulated in individuals with severe and symptomatic malaria (3928) were downregulated in those with asymptomatic malaria or individuals not infected. Consequently, genes that were upregulated in asymptomatic malaria infected individuals or those with no infection (1604) were downregulated in severe/asymptomatic malaria conditions. While 130 genes showed no change in the level of their expressions

**Figure 2:**
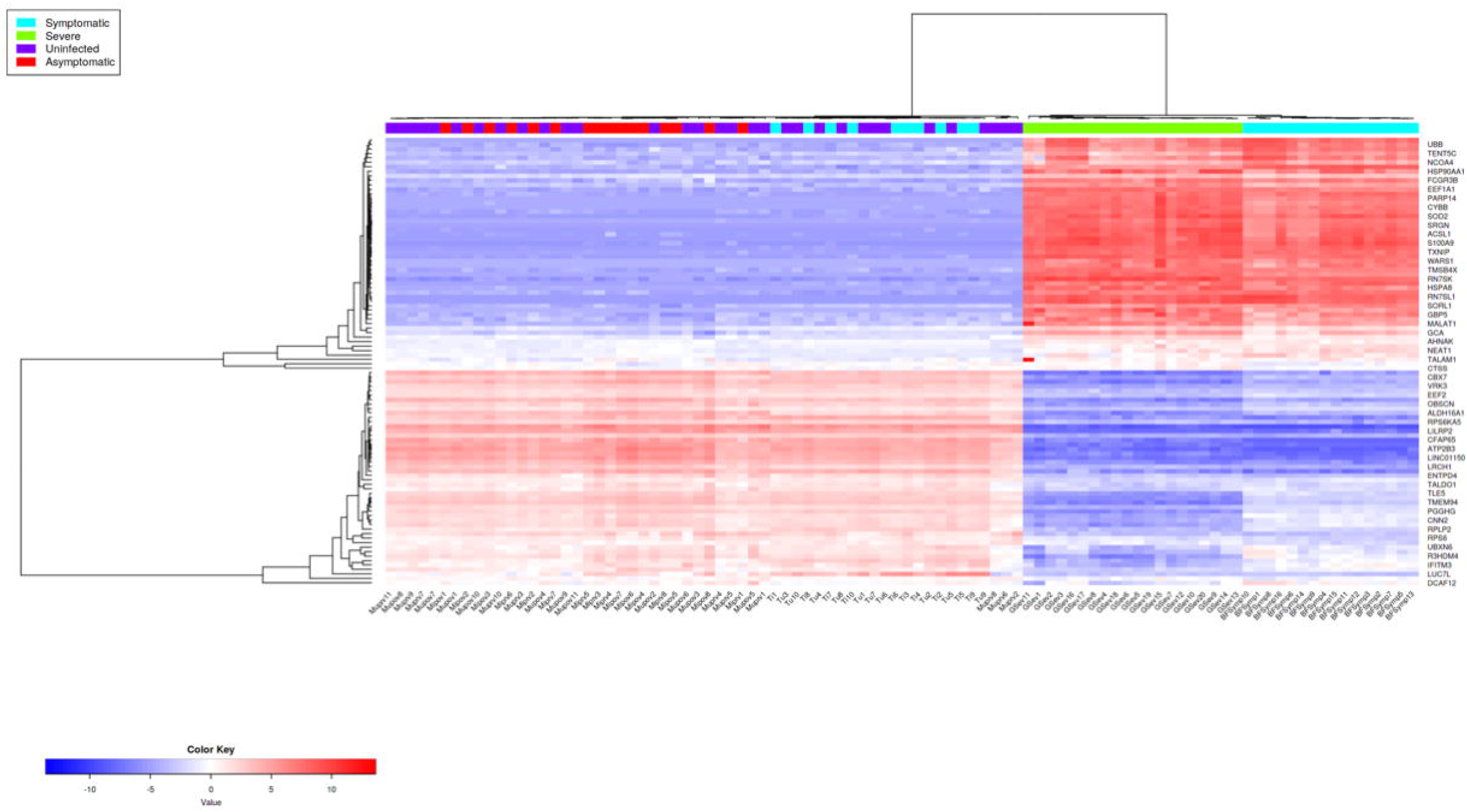
Differentially expressed genes in individuals with various *Plasmodium falciparum* clinical manifestations. Individuals with symptomatic and severe malaria showed similar patterns of up/downregulated genes, and this pattern was reversed in those with asymptomatic or no malaria infection. NB: GSev – Gambia severe; BFSymp – Burkina Faso Symptomatic; Tu – Tanzania uninfected; Ti – Tanzania infected; Muprv – Mali uninfected pre-vaccinated group; Mupov – Mali uninfected post-vaccinated; Miprv – Mali infected pre-vaccinated; Mipov – Mali infected post-vaccinated,

**Figure 3:**
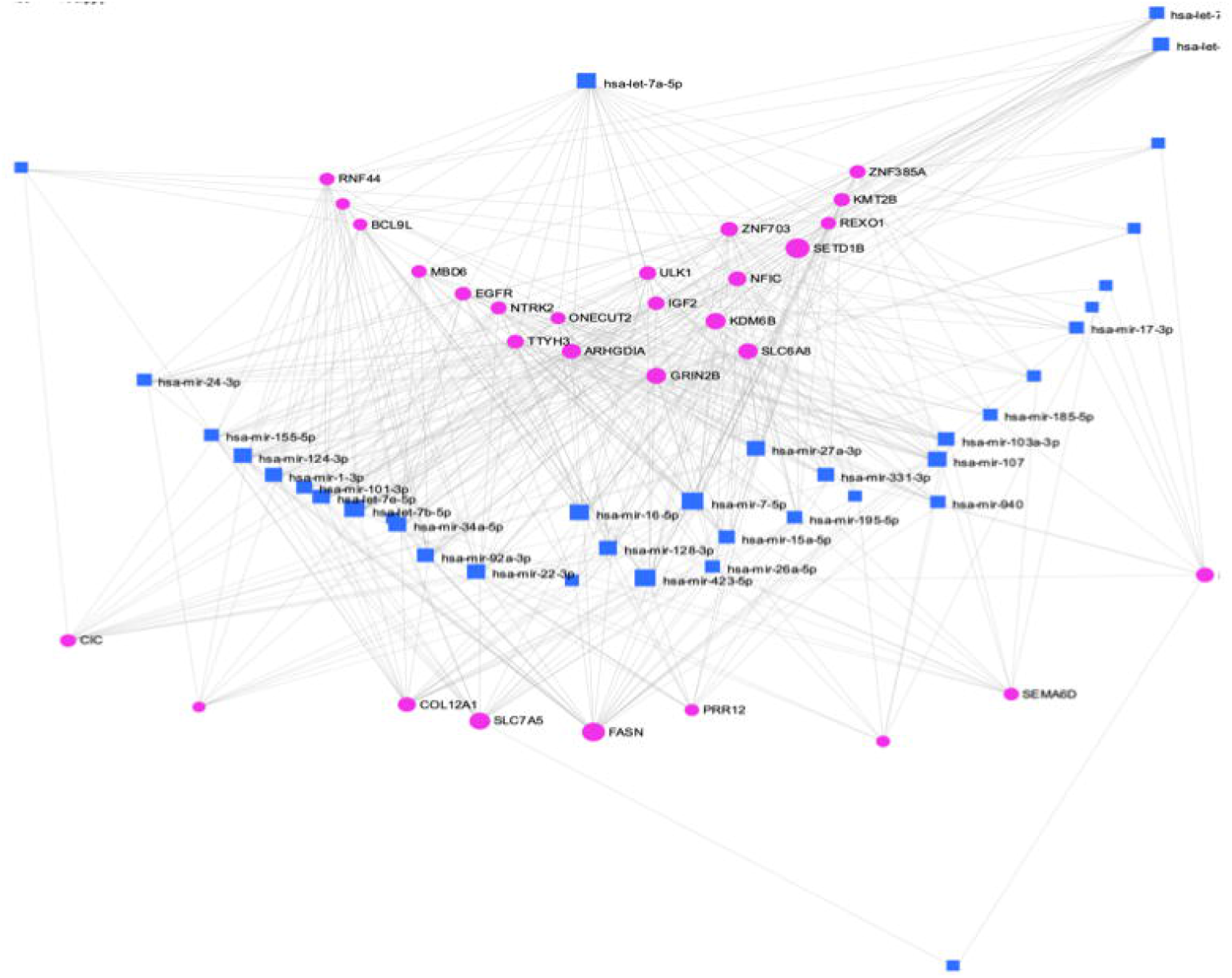
Network interactome between DEGs (pink) and differentially expressed miRNAs (blue). Among all the DEGs and miRNAs, these are the only ones that interacted with one another, indicating a pathway possibly implicated with malaria. Hsa-mir-331 interacting and regulating KMT2B, IDF2, TTYH3, and NTRK2 genes and hsa-mir-24 interacting with EGFR, MBD6, ZNF385A, as well as TTYH3 genes in this interactome. Among all the differentially expressed miRNAs, these are the ones interacting with the DEGs.

### Differentially expressed miRNA interacting with different target genes and lncRNAs in the dataset

A total of 141 miRNAs were differentially expressed. These miRNAs were found across all human chromosomes except chromosome 21, which had no representation. Of these, 78 were upregulated and 63 downregulated. Various biological processes including positive regulation of cell cycle (adjusted p-value 0.00000153), negative regulation of protein metabolism (adjusted p-value 0.0000194), response to drug (adjusted p-value 0.00197) were identified (Figure 4a) in the upregulated miRNA. Biological processes identified among the downregulated miRNA include regulation of RNA metabolic process (adjusted p-value 0.000144), regulation of gene expression (adjusted p-value 0.000201), regulation of transcription (adjusted p-value 0.000393), and cellular response to extracellular stimulus (adjusted p-value 0.000951) (Figure 5a). Molecular functions of upregulated miRNAs identified 25 functions including enzyme binding (adjusted p-value 0.0454), protein complex binding (adjusted p-value 0.0454) etc (Figure 4b). While 20 molecular functions including neutral amino acid transmembrane transporter activity (adjusted p-value 0.00527) and RNA binding (adjusted p-value 0.0408) (Figure 5b) were identified among the downregulated miRNAs. Of the 16 cellular components identified in miRNA that were overexpressed, cytosol (adjusted p-value 0.000179), mitochondrial outer membrane (adjusted p-value 0.000837), nucleoplasm (adjusted p-value 0.000895) top the list (Figure 4c), while ribonucleoprotein complex (adjusted p-value 0.0000549), nucleus (adjusted p-value 0.0309), spliceosome complex (adjusted p-value 0.0467) were among the cellular components observed in downregulated miRNAs (Figure 5c).

**Figure 4:**
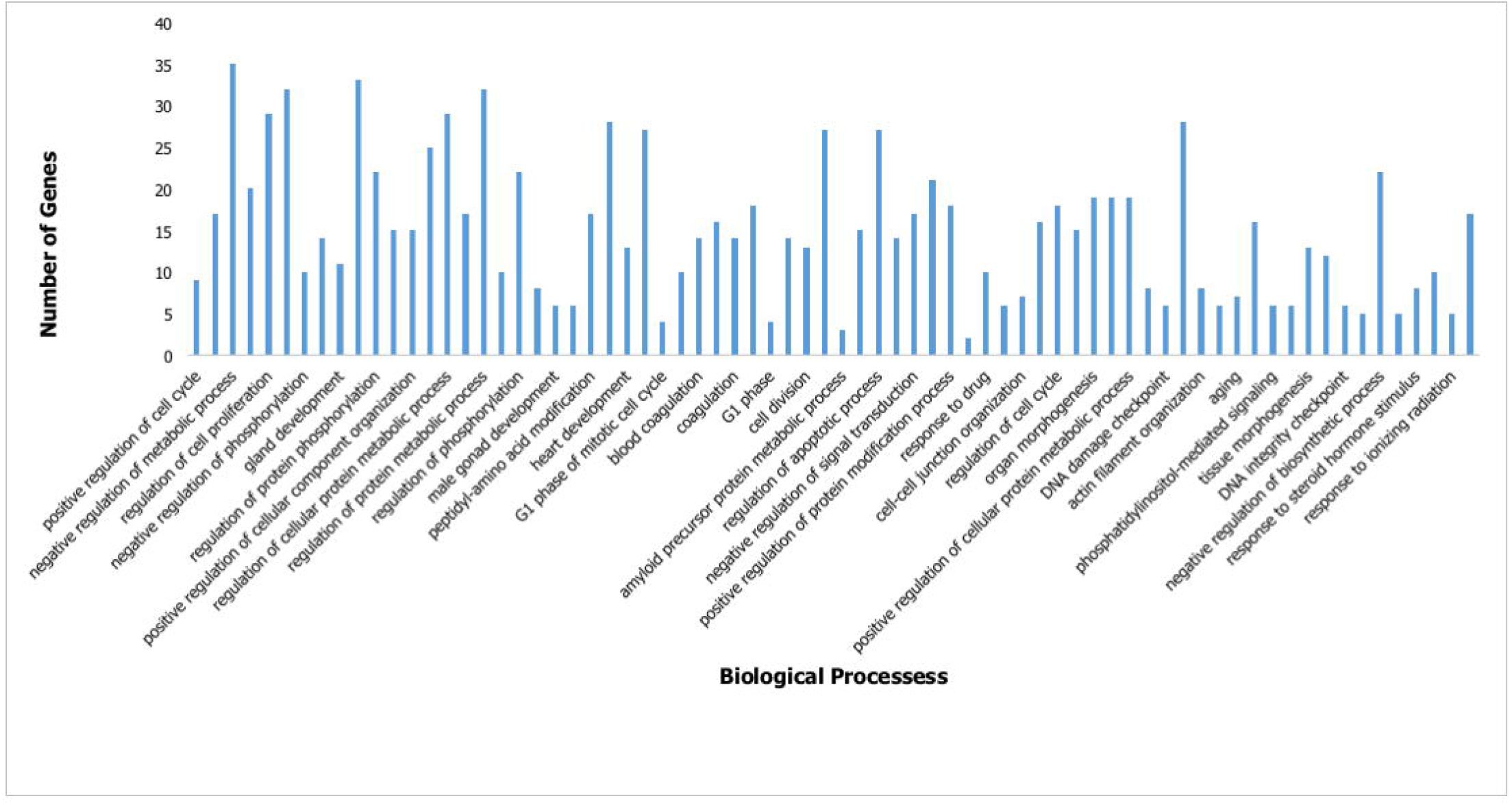
Enriched biological processes (a), molecular function (b) and cellular component (c) of upregulated miRNAs from individuals with different malaria clinical outcomes

**Figure 5:**
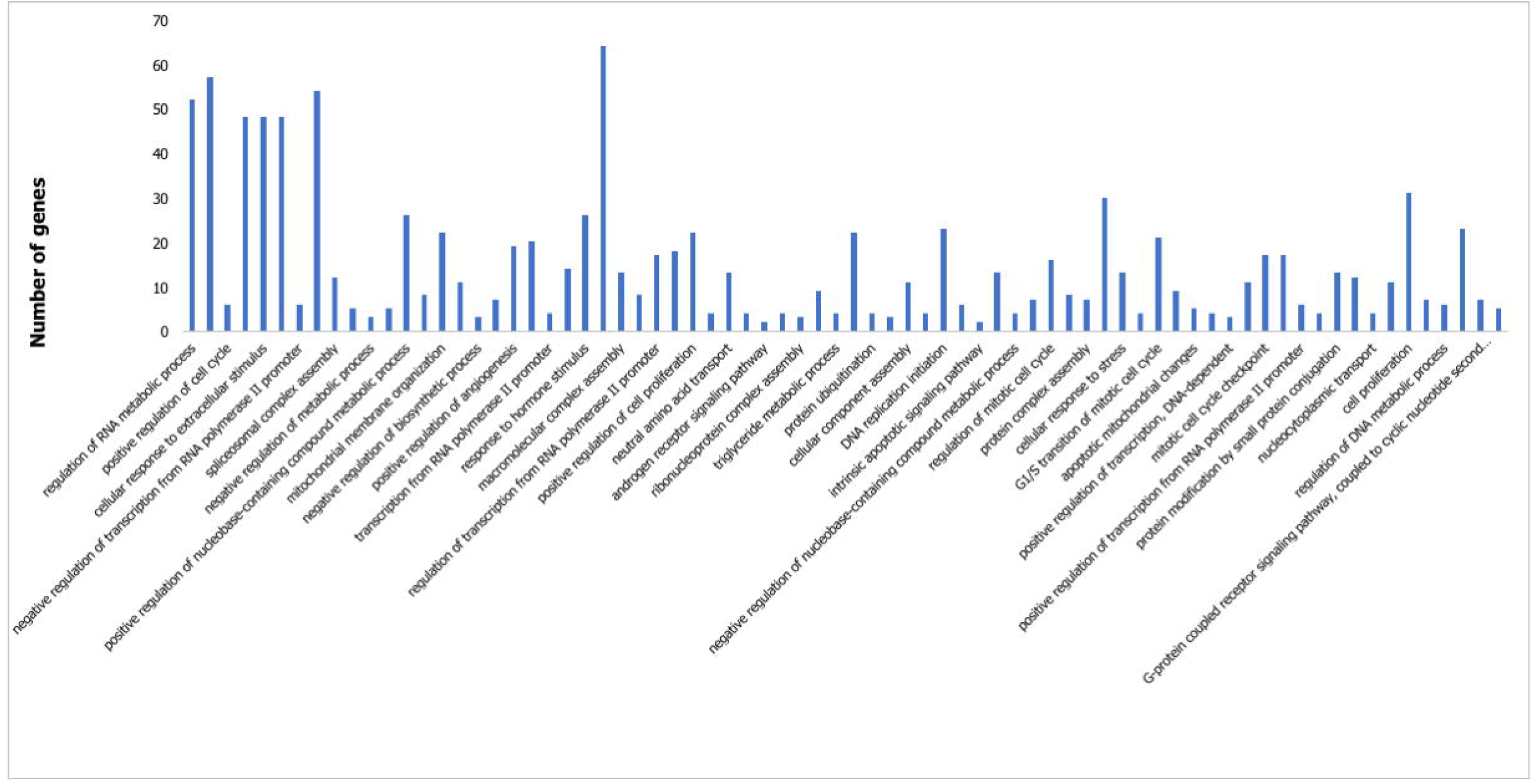
Enriched biological processes (a), molecular function (b) and cellular component (c) of downregulated miRNAs from individuals with different malaria clinical outcomes

Of all identified miRNA, five stood out (hsa-mir-32, hsa-mir-25, hsa-mir-221, hsa-mir-29 and hsa-mir-148), and were shown found to have multiple interactions with various genes. Of these five, hsa-mir-221 had the highest connections to 16 genes, including DDIT4, ICAM1, CDKN1C, p27, FOS etc. The second most connected miRNA is hsa-mir-29a with connections to 14 genes (CDK6, CD276, CDC42, PTEN, DNMT3A, CXXC6 etc), while the least connected miRNA is hsa-mir-32 with only one interaction (Figure 6). Furthermore, all of the five miRNAs, with the exception of hsa-mir-32, interacted with multiple lncRNAs, such as hsa-mir-221 interacting with RUNDC3A, TMEM147, FGDS, ARHGA27P1, hsa-mir-29a with STAG3L5P, MALAT1, hsa-mir-25 with NEAT1, LINC02275, and hsa-mir-148 with MMP25, XIST, ERVK13 (Figure 7).

**Figure 6:**
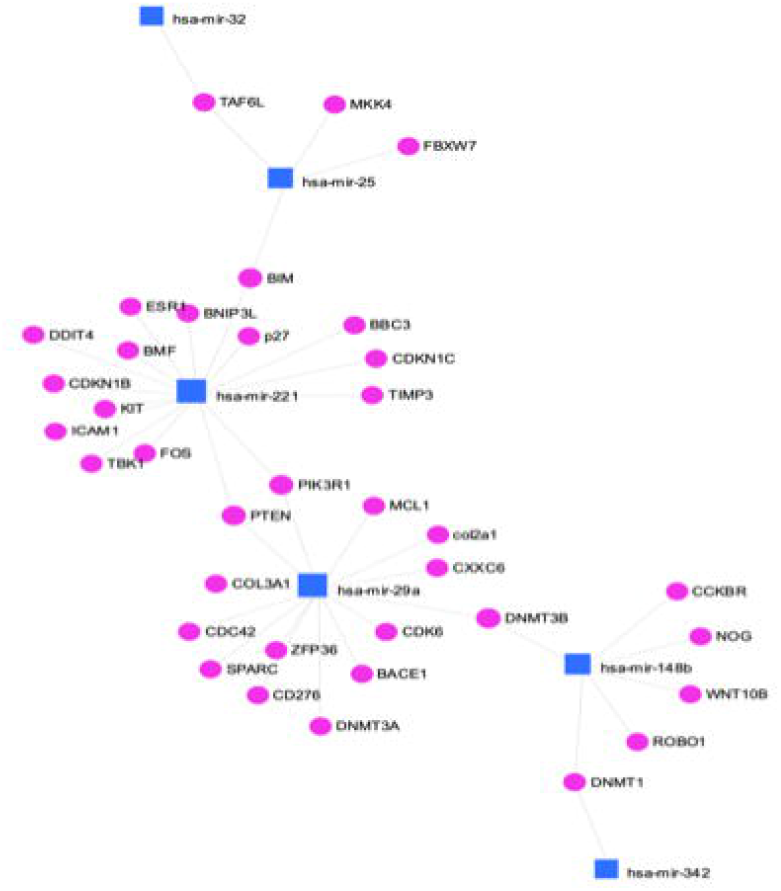
Network interaction between the most unique miRNAs (blue) and target genes (pink). Hsa-mir-221 (16 genes), hsa-mir-29a (13 genes) and hsa-mir-148b (6 genes) had the most connections, while hsa-mir-342 and hsa-mir-32, were the least connected (one gene each). It is postulated that genes with the most interactions could serve for diagnostic or therapeutic purposes in managing disease.

**Figure 7:**
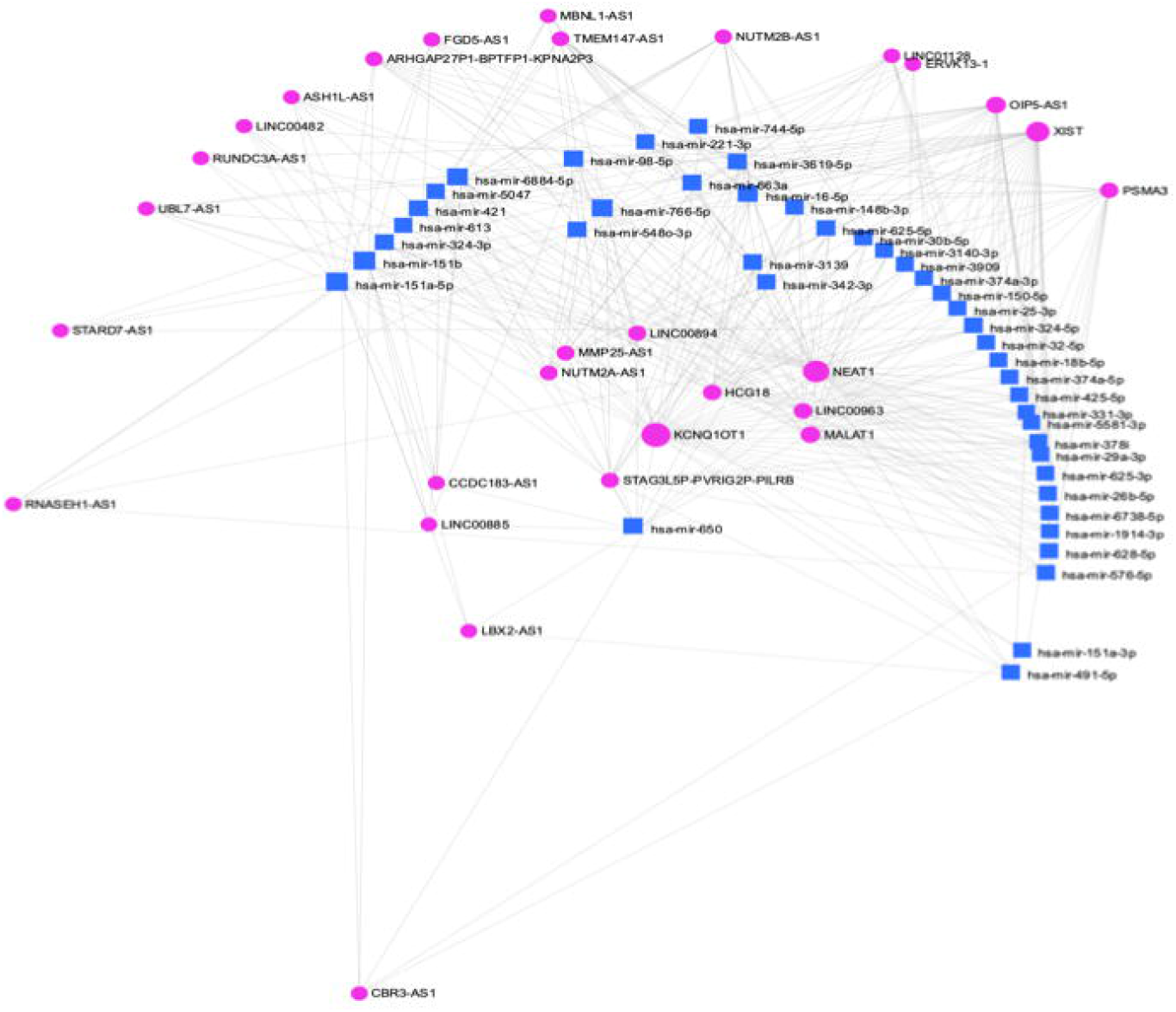
Network interaction between predominant miRNAs (blue) and some lncRNAs (pink). Various interactions was observed in this visualized display.

### Characterization of identified lncRNA

A total of 608 lncRNA were identified to have various expression patterns, with 586 and 22 recorded to be upregulated and downregulated respectively. Applying further stringency reveal only 522 and 14 lncRNAs to be upregulated and downregulated respectively (Supplementary Table 1 and 2). The minimum number of exons found among the highly expressed lncRNA is 2, while the maximum is 32, however, among the downregulated lncRNAs, the minimum exon is 2 and the maximum 6. Some of highly expressed lncRNAs such as SLC7A11 through interactions with genes like CD43, CDK41, ZBTB47 were found to be involved in antigen-specific activation of T cells, regulation of transcription by RNA polymerase II, while LINC01524 is involved in CD28 co-stimulation when it interacts with CD292, IRGQ, HOOK3, BRD4, AP1M1 genes (Supplementary Table 1 and 2).

### Biological, molecular and cellular functions of the differentially expressed genes

The pathway analyses of the top 100 upregulated genes identified 68 biological processes (Figure 8a), including axon guidance (p-value 2.52E-07), cell-cell adhesion (p-value 3.78E-05), transmembrane receptor protein tyrosine kinase signaling pathway (p-value 4.33E05), and hemophilic cell adhesion via plasma membrane adhesion molecules (p-value 1.87E040). On the other hand, enrichment analyses of the top 100 downregulated genes identified 26 biological processes (Figure 8b), including oxygen transport (p-value 6.64E-09), hydrogen peroxidase catabolic process (p-value 1.43E-07), cellular oxidant detoxification (p-value 2.42E-05) the most enriched.

**Figure 8:**
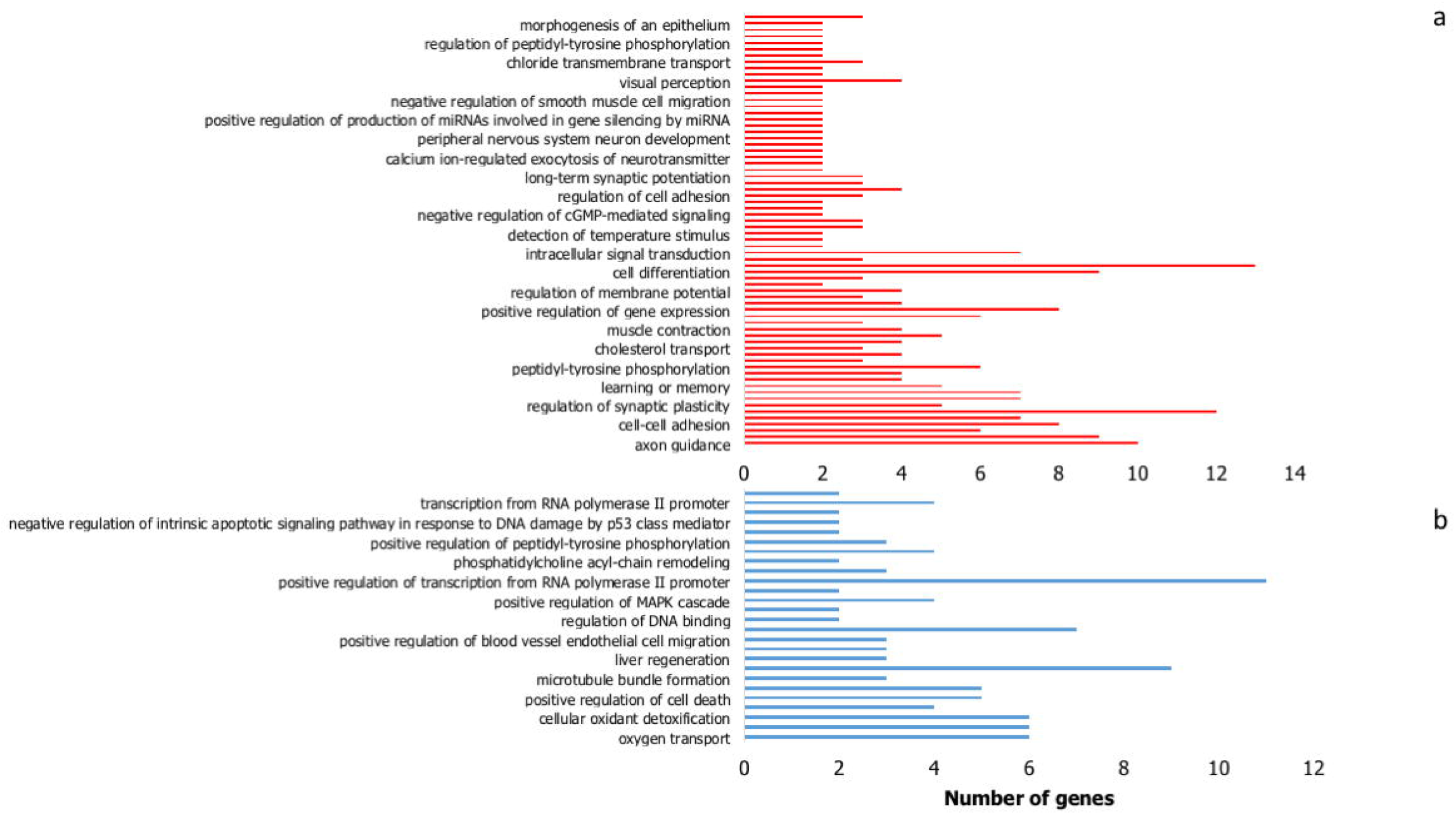
Biological processes identified among the topmost 100 (a) upregulated and (b) downregulated genes from individuals with different clinical malaria manifestation

Thirty-six (36) molecular functions were identified from these upregulated genes, with functions such as extracellular matrix constituent lubricant activity (p-value 2.34E04), extracellular matrix structural constituent (p-value 5.83E04), protein binding involved in cell-cell adhesion (p-value 0.00823392) were among the most significant pathways (Figure 9a). However, only 16 molecular functions were identified in the probed downregulated genes, of which oxygen transport activity (p-value 9.54E-11), haptoglobin binding (p-value 6.55E-10) and organic acid binding (p-value 1.20E-09) were more enriched than others (Figure 9b).

**Figure 9.**
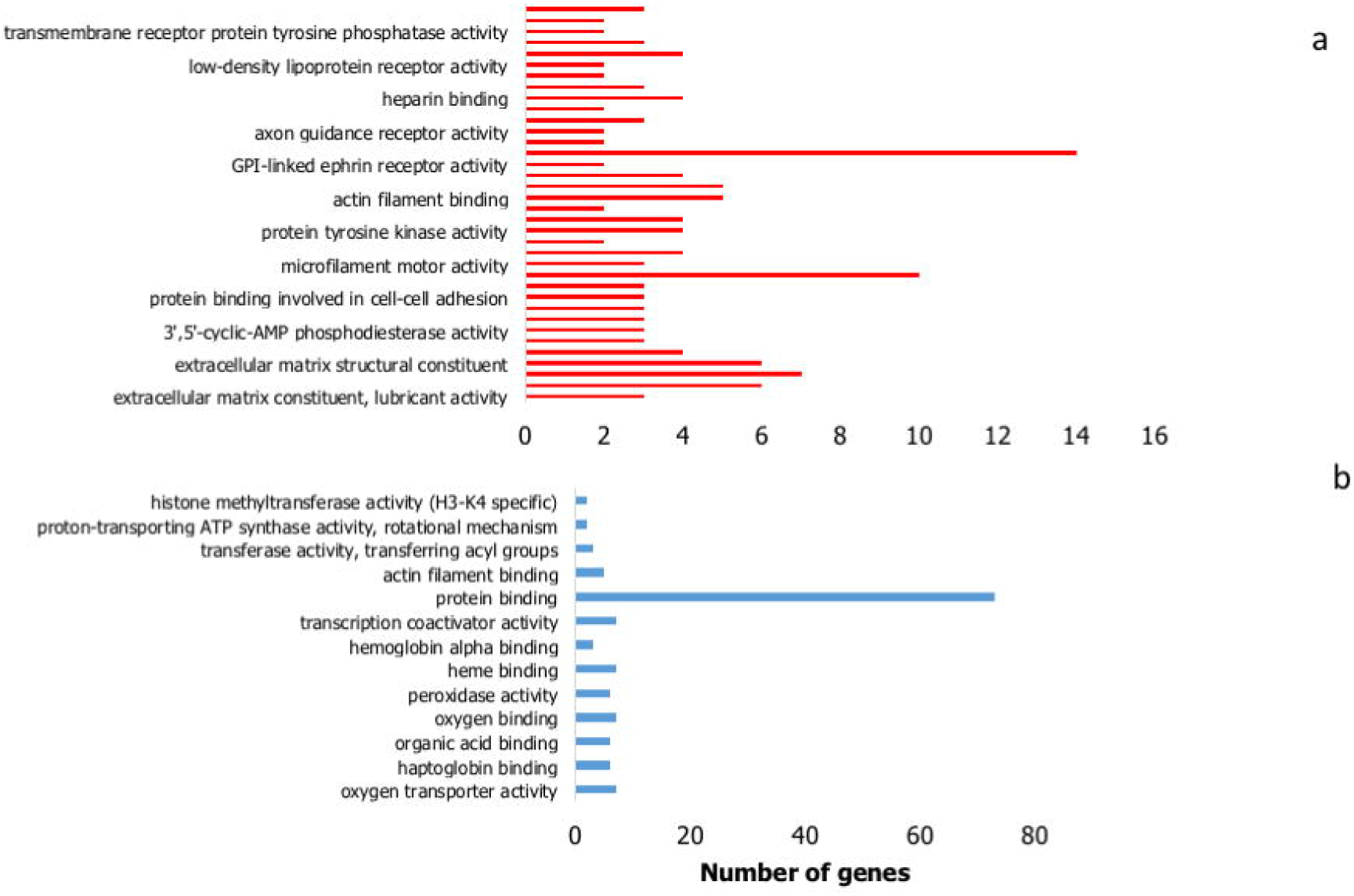
Molecular Functions identified among the topmost 100 (a) upregulated and (b) downregulated genes from individuals with different clinical malaria manifestation

Integral component membrane (p-value 5.65E-08), plasma membrane (1.79E-07), receptor complex (p-value9.95E-07) were the most enriched cellular components of the upregulated genes (Figure 10a), while only 11 cellular components were identified among the downregulated ones, with hemoglobin complex (p-value 2.32E-11) and haptoglobin-hemoglobin complex (p-value 8.28E-10) the most enriched (Figure 10b). KEGG pathway analysis identified axon guidance, protein digestion and absorption, morphine addiction, and calcium oxytocin signaling pathways among the upregulated genes while MAPK signaling pathway, amyotrophic lateral sclerosis and tuberculosis the most identified among the downregulated genes (Figure 11).

**Figure 10.**
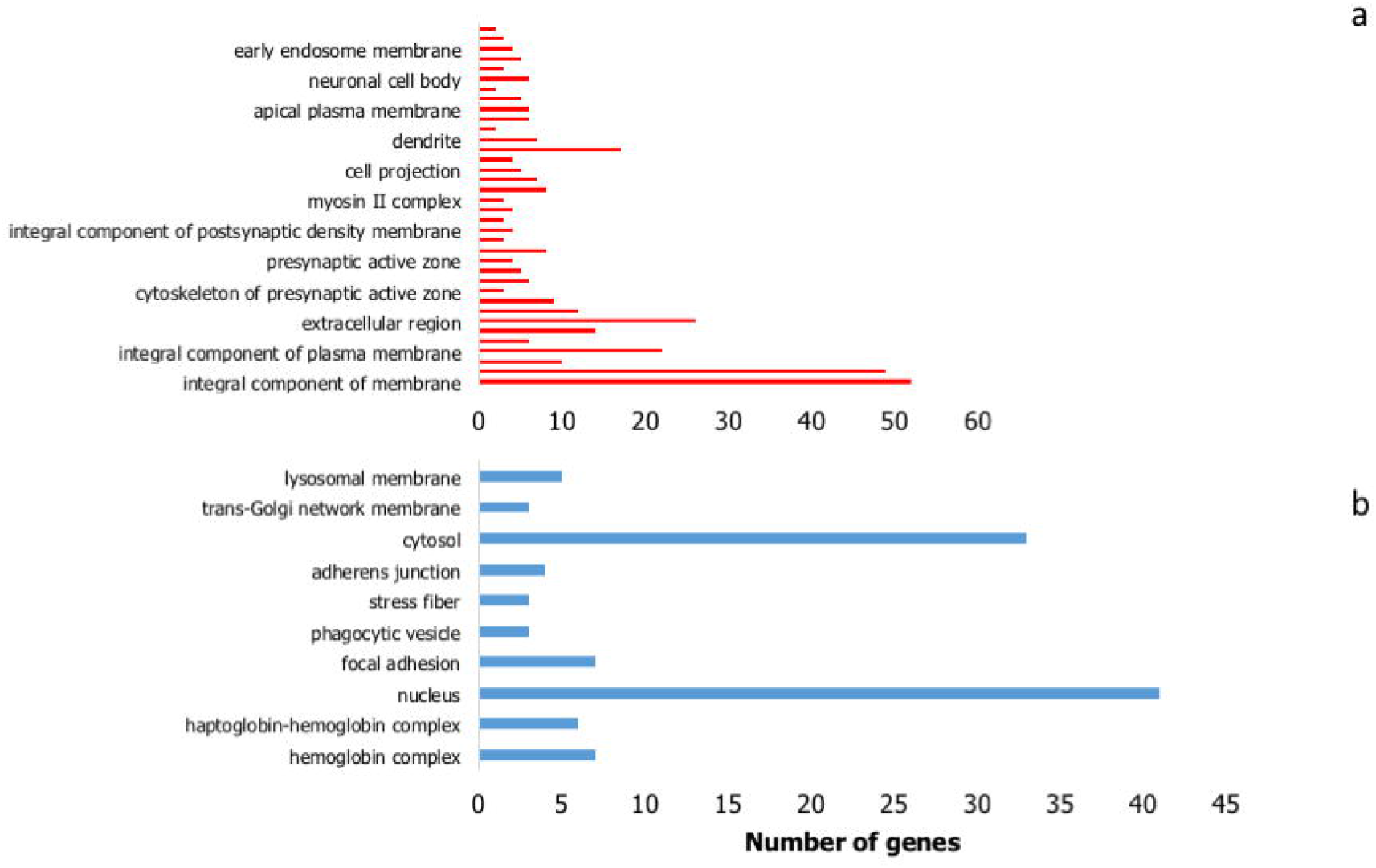
Cellular components identified among the topmost 100 (a) upregulated and (b) downregulated genes from individuals with different clinical malaria manifestation

**Figure 11.**
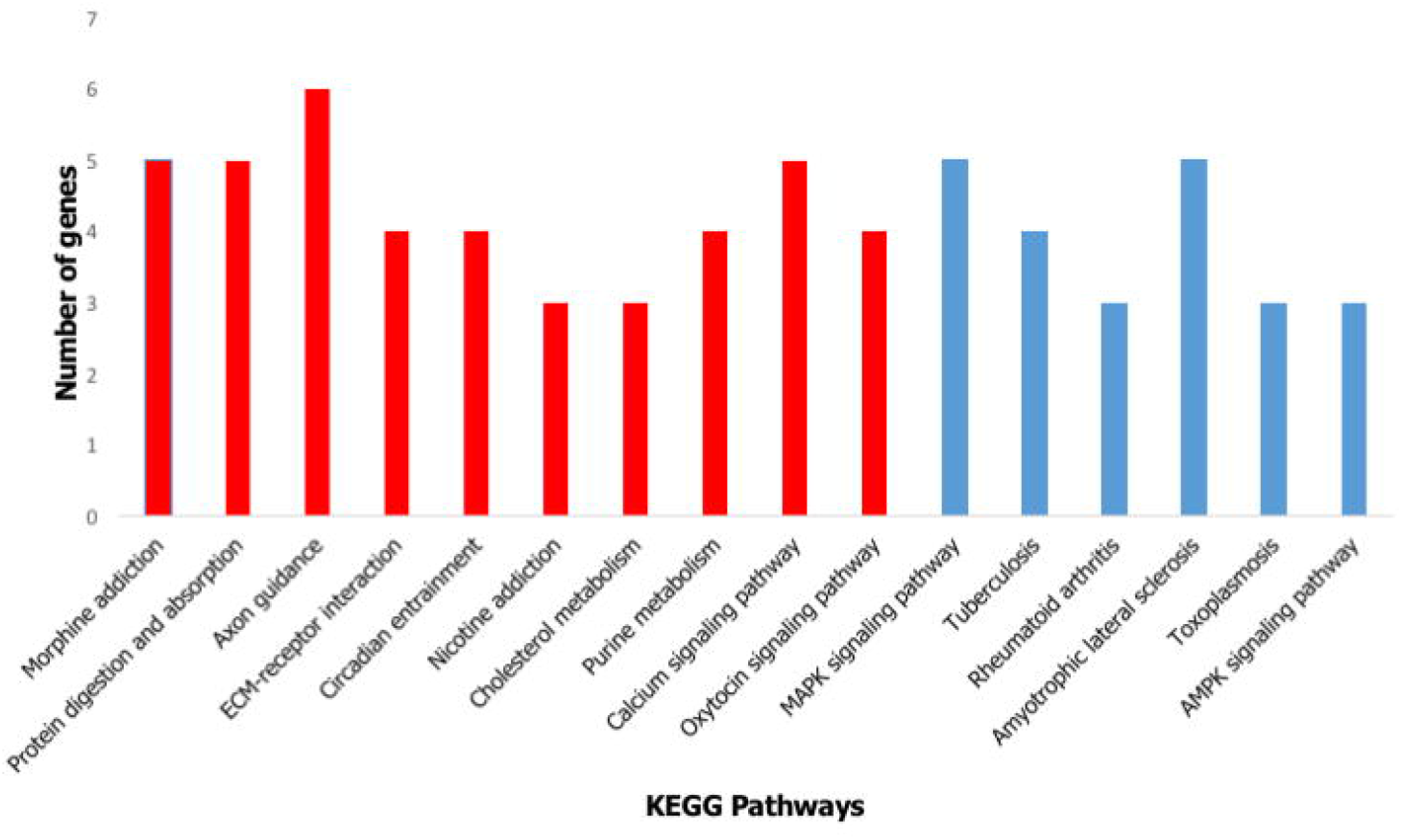
KEGG pathway analysis of upregulated genes (red) and downregulated genes (blue)

## Discussion

*P. falciparum* infection remains a major public health challenge in many countries, especially in sub-Saharan Africa, where children under five years of age and pregnant women are the most vulnerable and affected of the population, with infection leading to severe complications and outcome, if not promptly diagnosed and treated. Understanding how immune response genes and regulatory factors impact clinical outcome in different disease classifications can potentially unlock the key for new diagnostic biomarkers and druggable targets.

We used a computational approach to elucidate differentially expressed genes and multi-omic regulatory factors (miRNA and lncRNA) that are potentially contributory to disease outcome in different types of clinical malaria from the perspective of the human host. We identified large number of genes that were differentially expressed in the four clinical categories. Interestingly, the severe and symptomatic disease categories showed similar expression patterns, with same genes upregulated and downregulated in both groups. Among the upregulated genes, C-C motif chemokine 11 (CCL11), one of chemokine genes clustered around chromosome 17 and displaying chemotactic activity for eosinophils, as well as a participant in innate immune response (Nazarinia et al., 2022), was the most upregulated. It is predicted to induce the production of reactive oxygen species (ROS) in microglia cells and mediating neutrophils homing into damaged or infected areas, amplifying inflammatory response in the process (Adzemovic et al., 2012). Possibly of most importance today, elevated plasma levels of CCL11 has been associated with severe COVID-19 cases in hospitalized patients (Nazarinia et al., 2022). Another highly expressed gene observed is the mitochondrial RNR2L10 (mt-rnr2l10) gene, which is involved in the regulation and execution of apoptotic cell death during inflammation (Guo et al., 2003). Hence, we can surmise that these highly expressed genes in severe and symptomatic malaria individuals are significant contributors to the inflammatory process mediating a robust innate response, underpinned by an effective downstream adaptive immune response to disease, and ultimately clinical outcomes.

The gamma globin gene 2 (HBG2) among others, was found to be downregulated. This gene alongside HBG1 are normally expressed in fetal liver, bone marrow and spleen and replaced by adult hemoglobin (HbA) after birth (Pereira et al., 2015). Being linked to iron and oxygen binding, it is important for effective transportation of oxygen via heme, thereby preventing and reducing cell death. But in downregulated disease conditions as observed, cell death becomes inevitable, hence the severity of disease. In this study, we did not identify the C6KTB7 or C6KTD2 identified by others (Agamah et al., 2021) to be to be differentially expressed in malaria patients. A possible reason for this could be difference in the demographics of participants from whom the data were obtained as signatures of immune response is is shaped partly by the array of pathogens in different geography that challenge individuals residing in those areas.

Of interest is the identification of several miRNAs that were either upregulated or downregulated in our study. Five of the miRNAs - hsa-mir-32, hsa-mir-25, hsa-mir-221, hsa-mir-29 and hsa-mir-148 were peculiar in that they bind to different target genes and the possibility of regulating their expression during infection. To the best of our knowledge, this is possibly the first report identifying these five miRNAs as differentially expressed and regulating different host genes such as CDK6, CD276, CDC42, DNMT3A, CXXC6 etc. CDK6 is known to phosphorylate and regulate the expression of tumor suppressor proteins, and its altered expression has been observed in prostate cancer (Chen et al., 2022), stomach cancer (Liu et al., 2022), and coronary artery disease (Witten et al., 2022). CD276 belongs to the immunoglobulin superfamily of proteins and participate in the regulation of T-cell-mediated immune response (Liu et al., 2021). It is highly expressed, hence of no surprise that hsa-mir-29 binds to the 3’UTR resulting in even higher expression that ultimately leads to inflammatory response and cell death. However, a study has found an inverse relationship between hsa-mir-29 and the expression of cytochrome P450 2C19 (CYP2C19) gene (Yua et al., 2015), where hsa-mir-29 was reported to suppress the expression of CYP2C19 gene.

Substantial number of lncRNAs were also identified to be associated with disease. Of these, two (SLC7A11 and LINC01524) were found to interact and stimulate several immune genes such as CD43, CDK41, ZBTB47, CD292, IRGI etc which are known to be involved in antigen-specific activation of T cells and regulation of transcription.

Our pathway analyses of differentially expressed genes reveal several pathways such as cell-cell adhesion, transmembrane receptor protein tyrosine kinase signaling pathway, hemophilic cell adhesion, protein binding, oxygen transport, hydrogen peroxidase catabolic process, cellular oxidant detoxification, hemoglobin complex and haptoglobin-hemoglobin complex that were enriched among the up-/downregulated genes. Multiple genes were identified to be involved in in the different pathways. For example, EGFR, EPHAS, and ROS1 genes implicated in lung cancer susceptibility (Ke et al., 2018; Zhang et al., 2022; Zhu et al., 2016) and were involved in positive regulation of kinase activity, cell-cell adhesion etc.

Several upregulated and downregulated lncRNAs were identified. Of these, SLC7A11, LINC02275, ERICH3, LINC01120, FOXD2, LINC01816 were unique in the number of target immune genes they bind from CD43, TNFRSF13B to CD292. Therefore, we proposed that these lncRNAs could that have the ability to regulate the expression of these various genes and impact their over-or under expression can be targeted as biomarkers for diagnostic purposes in distinguishing the different forms of malaria infections.

Evaluating the network connection between the DEGs and miRNA, differentially expressed miRNAs and various genes and miRNAs and lncRNAs to appreciate the visual connection between all revealed various interesting networks amongst the duplexes, for instance interaction was observed between KMT2B/IDF2/TTYH3/NTRK2 genes and hsa-mir-331, EGFR/MBD6/ZNF385A/TTYH3 and hsa-mir-24. The most connected miRNA is the hsa-mir-221 which was observed to target 16 genes including ICAM1, followed by the hsa-mir-29a binding to 14 genes (including CD276, CDC42, and CXXC6). Similarly, these two miRNAs were also found to be connected to several lncRNAs. Taken together, it is safe to propose that these two miRNAs play significant role in human immune response to malaria and results in different clinical outcomes. Therefore, since they modify and/or regulate the expression of these genes and lncRNAs, targeting them for diagnostic purpose in discriminating individuals with severe/symptomatic malaria from those with asymptomatic infection or not infected will provide an added arsenal in the battle against malaria.

This is a computational study utilizing RNAseq transcriptome data to decipher DEGS, miRNAs and lncRNAs involved in malaria infection and how they modulate disease status and severity (outcome). Although the results from our analysis could not be validated with clinical cases, we strongly opine that the findings can serve as the basis for subsequent *in vitro* and in vivo characterization of individuals with different malaria clinical outcomes.

## Acknowledgment

We acknowledge ongoing support and funding from the College of Health Sciences and Technology, Rochester Institute of Technology (BNT). MAO is supported through the American Association of Immunologists Intersect Fellowship Program for Computational Scientists and Immunologists. The funders had no role in study design, data collection and analysis, decision to publish, or preparation of the manuscript.

## Author contributions

BNT conceptualized this project; MAO and OBM carried out initial *in silico* survey; MAO performed the experiment, analyzed the data and wrote the first draft of the manuscript; OBM, OO and BNT contributed to the discussion and revision of the manuscript. All authors reviewed and approved the final manuscript.

